# NATURAL ENEMIES OF HERBIVORES MAINTAIN THEIR BIOLOGICAL CONTROL POTENTIAL UNDER FUTURE CO_2_, TEMPERATURE AND PRECIPITATION PATTERNS

**DOI:** 10.1101/2020.07.15.204503

**Authors:** Cong Van Doan, Marc Pfander, Anouk Guyer, Xi Zhang, Corina Maurer, Christelle A.M. Robert

**Affiliations:** Institute of Plant Sciences, University of Bern, Altenbergrain 21, 3013 Bern, Switzerland; Oeschger Centre for Climate Change Research (OCCR), University of Bern, Falkenplatz 16, 3012 Bern, Switzerland; Agroscope, Müller-Thurgau-Strasse 29, 8820 Wädenswil, Switzerland; Key Laboratory of Plant Stress Biology, State Key Laboratory of Cotton Biology, School of Life Sciences, Henan University, Kaifeng 475004, China; Agroecology and Environment, Agroscope, Reckenholzstrasse 191, 8046 Zürich, Switzerland

**Keywords:** Climate change, Trophic interactions, Herbivore natural enemies, Spiders, Ladybugs, Wasps, Nematodes

## Abstract

Climate change will profoundly alter the physiology and ecology of plants, insect herbivores and their natural enemies, resulting in strong effects on multitrophic interactions. Yet, manipulative studies that investigate the direct combined impacts of changes in CO_2_, temperature, and precipitation on this group of organisms remain rare. Here, we assessed how three day exposure to elevated CO_2_, increased temperature, and decreased precipitation affect the performance and predation success on species from four major groups of natural enemies of insect herbivores: an entomopathogenic nematode, a wolf spider, a ladybug and a parasitoid wasp. Future climatic conditions (RCP 8.5), entailing a 28% decrease in precipitation, a 3.4°C raise in temperature and a 400 ppm increase in CO_2_ levels, slightly reduced the survival of entomopathogenic nematodes, but had no effect on the survival of other species. Predation success was not negatively affected in any of the tested species, but was even increased for wolf spiders and entomopathogenic nematodes. Factorial manipulation of climate variables revealed a positive effect of reduced soil moisture on nematode infectivity, but not of increased temperature or elevated CO_2_. These results suggest that natural enemies of herbivores are well adapted to short term changes in climatic conditions and may not suffer from direct negative effects of future climates. These findings provide mechanistic insights that will inform future efforts to disentangle the complex interplay of biotic and abiotic factors that drive climate-dependent changes in multitrophic interaction networks.

## INTRODUCTION

Climate change entails increasing concentrations of greenhouse gases in the atmosphere, rising temperatures, and shifts in precipitation patterns (IPCC 2014). These changes will profoundly alter individual species’ physiologies and ecology, interaction networks, community composition and (agro)ecosystem functioning (Bellard *et al*. 2012; Walther 2010; Boukal *et al*. 2019; Parmesan & Yohe 2003; Newman *et al*. 2011; Abdala-Roberts *et al*. 2019; Selvaraj *et al*. 2013). The feeding efficiency of natural enemies of herbivores in particular was predicted to be altered by climate change (Tylianakis *et al*. 2008; Voigt *et al*. 2003; Rosenblatt & Schmitz 2016), which may lead to strong effects on trophic interaction networks and biological control services (Vidal & Murphy 2018; Gravel *et al*. 2016; Woodward & Bohan 2013; Bohan *et al*. 2013; van der Putten *et al*. 2004; Altieri *et al*. 2012). Yet, manipulative studies that investigate the combined impacts of changes in CO_2_, temperature, and precipitation on members of higher trophic levels remain rare (Jamieson *et al*. 2012; Rosenblatt & Schmitz 2016; Laws 2017), thus limiting our capacity to predict how climate change will affect natural enemies of herbivores.

Natural enemies of herbivores such as predators, parasites and parasitoids may suffer from climate change via extrinsic and intrinsic mechanisms. Intrinsic mechanisms refer to direct effects of climate-associated abiotic parameters on an organism (Cornelissen 2011; Robinet & Roques 2010). Extrinsic mechanisms refer to indirect effects, mediated for instance through shifted distributions, altered phenology or physiology of lower trophic levels (Han *et al*. 2019; Kaplan *et al*. 2016; Kharouba *et al*. 2018; Jeffs & Lewis 2013; Pincebourde *et al*. 2017; Chidawanyika *et al*. 2019; Damien & Tougeron 2019; Renner & Zohner 2018). While investigating indirect effects of climate change across trophic levels is crucial to predict community shifts (van der Putten *et al*. 2010), understanding the direct impact of climate change onto each trophic level is essential to characterize the underlying mechanisms (Thomson *et al*. 2010).

Over the last years, detailed experiments have been conducted to evaluate the direct impact of individual climate-change associated abiotic factors on natural enemies of herbivores. For example, temperature was identified as a key determinant of parasitoid developmental rates, survival, fecundity, parasitism and dispersal (Hance *et al*. 2007; Walther *et al*. 2002; Selvaraj *et al*. 2013). Thermal performance curves revealed that increasing temperatures enhance parasitism success until a maximum at optimum temperature, beyond which any increase in temperature leads to a decline of parasitism success (Furlong & Zalucki 2017; Chidawanyika *et al*. 2019). Optimum temperature was repeatedly reported to be lower for parasitoids than for their hosts, indicating that parasitoids may be more susceptible to elevated temperature than herbivores, resulting in lowered biocontrol efficacy over time (Furlong & Zalucki 2017). Similarly, changes in CO_2_ levels and precipitation were demonstrated to affect natural enemies of herbivores. While the impact of elevated CO_2_ on the third level are generally mediated by shifts in bottom-up forces, direct effects of CO_2_ on predators and parasitoids should not be neglected (Jun Chen *et al*. 2007). Notably, elevated CO_2_ alters the chemosensation of natural enemies by disrupting their ability to perceive or process cues from their environment (Draper & Weissburg 2019), for instance through a reduction of olfactory receptor sensitivity and habituation of higher-order neurons (Majeed *et al*. 2014). Changes in precipitation were reported to disturb the physiology and behaviour of natural enemies of herbivores (Torode *et al*. 2016; Jamieson *et al*. 2012; Barnett & Facey 2016). While arachnids and insects possessing a waxy cuticle can reduce evaporation (Berridge 2012), soft body organisms (nematodes, isopods, and myriapods) may suffer larger water loss under drought conditions (Sylvain *et al*. 2014). Drought may also prompt arthropod predators to migrate, hide in the soil or build shelters to avoid desiccation (Willmer 1982; Berridge 2012; Melguizo-Ruiz *et al*. 2016), thus reducing their foraging activity.

Compared to single climate parameters, much less is known about the direct effects of combined and interactive impact of multiple parameters on natural enemies of herbivores (Robinson *et al*. 2012; Kreyling & Beier 2013; Jactel *et al*. 2019; Hiltpold *et al*. 2016). A meta-analysis on the impact of multiple global change drivers on the survival of animals from all trophic levels revealed that combined climatic stressors led to non-additive effects, comprising synergistic and antagonist interactions, in the majority (65%) of the examined studies (Darling & Côté 2008). The occurence of synergistic and antagonist effects varies considerably between meta-analyses and is species and sex-dependent (Darling & Côté 2008; Crain *et al*. 2008; Rosenblatt & Schmitz 2014). Studies evaluating the direct effects of combined climatic factors on herbivore natural enemies remain scarce. One example showed antagonist interactions between elevated temperature and decreased precipitation pattern on parasitoid wasp success: while elevated temperature and drought individually enhanced parasitism success, their combination reversed this effect (Romo & Tylianakis 2013). Another study, using altitudinal gradient experiments, revealed that warming, elevated nitrogen resources and their combination resulted in similar increase in total herbivore biomass, but not that of parasitoids (Sassi & Tylianakis 2012). Even though parasitoids parasitized more herbivores under elevated temperature, this increase was not proportionate to the increase in host abundance. This observation could be explained by the negative effect of both drivers and their sub-additive interactions on parasitism rates (Sassi & Tylianakis 2012). Despite these examples, to date, the outcome of multiple global change factors on the third trophic level remains unpredictable and is likely to lead to “ecological surprises” (Darling & Côté 2008).

Here, we developed a microcosm climate system to quantify the direct impact of elevated temperature (incl. an appropriate diurnal rhytm) and CO_2_ as well as reduced precipitation, as predicted by the Representative Concentration Pathway 8.5 (RCP 8.5, (IPCC 2014)), on the survival and foraging efficiency of different natural enemies of herbivores. Our work reveals that short-term exposure to RCP 8.5 conditions has no negative effects on several different natural enemies of herbivores, suggesting that the third trophic level is resilient to direct abiotic effects imposed by climate change.

## MATERIALS AND METHODS

### Biological resources

#### Herbivores

Eggs of the cucumber banded beetle, *Diabrotica balteata*, LeConte (Coleoptera: Chrysomelidae), were kindly provided by Oliver Kindler (Syngenta, Stein, Switzerland). Hatching larvae were reared on freshly germinated maize seedlings (*var*. Akku, DSP, Delley, Switzerland) until use. Third instar larvae were used for all infectivity experiments. Eggs of Egyptian cotton leafworm, *Spodoptera littoralis* (Lepidoptera: Noctuidae), were provided by Ted Turlings (Laboratory of Fundamental Research in Fundamental and Applied Chemical Ecology, University of Neuchâtel, Neuchâtel, Switzerland) and reared on artificial diet until use. Larvae of the common fruit fly, *Drosophila melanogaster* (Diptera: Drosophilidae), were provided by Dirk Beuchle (Group Suter, Institute of Cell Biology, University of Bern, Bern, Switzerland). Individuals of the bird cherry-oat aphids, *Rhopalosiphum padi* (Hemiptera: Aphididae), were bought from Andermatt Biocontrol (Grossdietwil, Switzerland) and reared on barley until use. Individuals of the cabbage aphid, *Brevicoryne brassicae* (Hemiptera: Aphididae), were collected in a cabbage field in Zürich (47°22’17.40” N/ 8°67’50.18” E, Agroscope in Wädenswil, Zürich, Switzerland).

#### Predators and parasites

Ladybirds, *Adalia bipunctata* (Coleoptera: Coccinellidae) and parasitoid wasps, *Aphidius ervi* (Hymenoptera: Aphidiidae) were bought from Andermatt Biocontrol (Grossdietwil, Switzerland). Wolf spiders, *Alopecosa albofasciata* (Araneae: Lycosidae), were collected in the field (Coordinates: 46°97’49.2” N/ 7°43’77.4” E, Bremgarten bei Bern, Switzerland). Entomopathogenic nematodes (EPNs), *Heterorhabditis bacteriophora*, strains EN01, TT01, DE6, HU2 and PT1 were obtained from in-house colonies (Zhang *et al*. 2019). All EPN strains were reared in larvae of the wax moth, *Galleria mellonella* (Lepidoptera: Pyralidae) (Fischereibedarf, Bern, Switzerland), until use (McMullen II & Stock 2014). Infective juveniles (IJs) were collected in aqueous suspension from white traps (White 1927) in tap water (1 EPN/μL) at room temperature for 10 days before use.

#### Current and predicted climatic conditions

Current and predicted climatic conditions were calculated using climatic data from the Swiss Central Plateau (Average of summer conditions from 2004 to 2016, Oensingen, 47°17’11.1” N / 7°44’01.5” E, Switzerland), data were supported by MeteoSwiss (Federal Office of Meteorology and Climatology, Zürich, Switzerland), and predictions from the Representative Concentration Pathway 8.5 (RCP 8.5, Intergovernmental Panel on Climate Change (IPCC) report (IPCC 2014). RCP 8.5 corresponds to an extreme scenario in which CO_2_ emissions continue to rise throughout the 21^st^ century. Consequently, current and RCP 8.5 atmospheric CO_2_ concentrations were of 450 ppm (± 50ppm), and 850 ppm (± 50 ppm) respectively. Current and RCP 8.5 of soil temperatures were 19.6°C and 23°C respectively. Because daily temperature variation can affect insect performance and predator-prey interactions (Stoks et al 2017), current and RCP 8.5 soil temperatures followed a diurnal variation of 3.5°C (minimal temperature at 6 am and maximal temperature at 4 pm) and reached a maximum temperature of 21.4°C and 24.8°C respectively (Figure S1). Current and RCP 8.5 soil volumetric moisture levels were adjusted to 23% and 16.6% (corresponding to 28% less precipitation (Figure S2)).

### Microcosm systems

To manipulate CO_2_ levels, temperature, and moisture, we developed a microcosm system using dry-bath cyclers and a custom-made CO_2_-dosage system. Falcon tubes (50 mL, Falcon, Greiner Bio-One, Frickenhausen, Germany) were filled with 30 g dry (80°C for 48 hr), sieved (2 cm mesh) soil (40% sand, 35% silt, 25% clay; Landerde, Ricoter, Aarberg, Switzerland). The natural soil microbiota was re-implemented to the soil as previously described (Hu *et al*. 2018). All falcon tubes were placed in dry bath cyclers (Digital Heating Cooling Drybath, Thermo Scientific, Fisher Scientific AG, Reinach, Switzerland) equipped with heating blocks that can accommodated up to nine falcon tubes. A CO_2_ mixing and distribution system was designed to continuously mix CO_2_ ambient air, measure the CO_2_ concentration of the mixture, and distribute it to different channels. Mixing CO_2_ and air was achieved using an air compressor (Prematic AG, Affeltrangen, Switzerland) coupled to two mass-flow-controllers (for CO_2_: Bronkhorst El-Flow Select F-200CV (0.6 mL.min^−1^), Ruurlo, Netherlands; and for air: CKD FCM-0010AI (0-10 L.min^−1^), CKD Corporation, Aichi, 485-8551, Japan). Ambient air from outside the building was used for mixing, therefore no CO_2_ was added to mimic current conditions (=450 ppm ± 50 ppm). A concentration of 400 ppm CO_2_ (purity 100%, 54.6 L bottle, and pressure of output at 0.8 bars, Gümligen, Switzerland) was added to ambient air (=850 ppm ± 50 ppm) to reach expected RCP 8.5 scenarios. The resulting CO_2_: air mix was pushed through a filter of activated carbon (Camozzi, Warwickshire, United Kingdom) and split through valves (Needle Valve 2839-⅛, CKD, Aichi, 485-8551, Japan) into seven individual channels in a series. The first channel, referred thereafter as “CO_2_ measuring channel”, was connected to a CO_2_ sensor (Rotronic AG, Bassersdorf, Switzerland). The air flow circulated alternatively between the CO_2_ measuring channel (for 2 min) and experimental channels (for 2 min). The two minutes duration between experimental channels was sufficient to reach stable expected CO_2_ concentrations. In all assays, four experimental channels were used, alternating between ambient (channels 2 and 4) and CO_2_ enriched (channels 3 and 5) air. Therefore, the ambient or CO_2_-enriched air was distributed through all channels within 16 min. This cycle was repeated every 30 min (16 min air distribution followed by 14 min pause) over the course of the experiment. Each of the experimental channels had 12 outlets (One-Touch fittings-male Straight, Sang-A Pneumatic Co., Daegu, Korea). Polyurethane tubing (outer/inner diameter: 4/2.5 mm, length: 2 m, Sang-A Pneumatic Co., Daegu, Korea) was connected to the outlets and distributed the air to the Falcon tubes. The tubing was attached to the lids of the Falcon tubes using One-Touch fittings-male Elbow (Sang-A Pneumatic Co., Daegu, Korea). The flow rate sent through individual Falcon tubes was adjusted to 1 L.min^−1^. The outflow of the Falcon tubes was connected to a collection system, itself connected to the CO_2_ sensor to verify CO_2_ levels. The collected air was then released in the environment.

The temperature in the Falcon tubes was controlled through the dry-bath cyclers and followed a diurnal variation of 3.5°C. Soil temperatures used to mimic current conditions were of 17.8°C at 6 am, and gradually increased to reach 21.4°C at 4 pm (Figure S1), as reported for the Swiss Plateau over the past two decades (MeteoSwiss, Federal Office of Meteorology and Climatology, Zürich, Switzerland). The temperatures mimicking the RCP 8.5 scenario were set to 21.2°C at 6 am and progressively increased to reach 24.8°C at 4 pm (Figure S1).

The moisture present in the tubes was controlled by adding the soil leachates to the tubes once at the beginning of the experiment. The volume of water to add in the tubes was calculated based on the soil density of 1.2 g.cm^−3^. Current moisture levels (23% soil moisture) were achieved by adding 16.6% (v/v) microbiota extracts contained in tap water and 6.4% (v/v) additional tap water. Predicted moisture levels (RCP 8.5, 28% less precipitation, Figure S2) were achieved by adding 16.6% (v/v) microbiota extracts contained in tap water only. The temperatures were adjusted to the different scenarios over a six-hour adaptation period (Figure S1). The water loss over the run of the experiments was not significantly different between treatments.

### Effects of climate change on aboveground predator and parasite survival

To evaluate the direct impact of current and predicted climatic conditions on success of predators, we evaluated the survival and predation success, defined as foraging efficiency, of a wolf spider (*A. albofasciata*, n=25-30), a ladybug (*A. bipunctata*, n=17) and a parasitic wasp (*A. ervi*, n=5-6) after exposure to climatic conditions in the microcosms. Half of the individuals for each predator species were exposed to current conditions, while the other half was exposed to RCP 8.5 conditions. A piece of cotton soaked in 10% sucrose solution or a slice of apple was added to the tubes as a sugar source. After three days of incubation, all animals were exposed to ambient temperature levels for one more day. This resulted in exposure to current and predicted temperature for three days, and to current or predicted CO_2_ and precipitation levels for 4 days prior predation assays. All predation assays were performed under ambient conditions.

### Effects of climate change on aboveground predation and parasitism success

To assess the predation and parasitism success of the three aboveground arthropods, four independent assays were conducted as follows. In a first experiment, individual wolf spiders were exposed to current or predicted RCP 8.5 climatic conditions as described above. After exposure, each spider was placed in a solo cup (250 mL, Pack Markt Sabaratnam, Solothurn, Switzerland). Half of the cups contained ten second-instar *S. littoralis* caterpillar larvae, while the second half contained ten *D. melanogaster* flies (n=9-12). In a second assay, individual ladybugs were exposed to current or RCP 8.5 climatic conditions (n=9-14) and then placed in petri dishes (94 x 16 mm, Bio-One Petri Greiner, Huberlab, Switzerland) containing a piece of cotton soaked in a 10% sucrose solution and thirty aphids *R. padi*. In the third experiment, three ladybugs per tube were exposed to current or predicted RCP 8.5 conditions (n=5-6) and then placed in petri dishes containing a piece of cotton soaked in a 10% sucrose solution and thirty aphids *B. brassicae*. In these assays, the number of consumed preys was recorded 24 hr after introduction of the predator. In a fourth assay, nine to twelve parasitoid wasps were placed in each tube and exposed to either current or predicted RCP 8.5 conditions (n=5-6). The wasps were then placed into petri dishes containing a slice of apple and thirty aphids *B. brassicae*. The wasps and aphids were left together for five days before assessing the aphid parasitism rate.

### Climate change on belowground parasitism rates

To investigate direct effects of an extreme climate change scenario (RCP 8.5) on the ability of soil-dwelling parasites to infect an herbivore, five strains of the entomopathogenic nematode *H. bacteriophora* (strains EN01, TT01, DE6, HU2, PT1) were tested in the microcosms described above. Briefly, 3’000 infective juvenile nematodes, suspended in one mL tap water, were added in each microcosm, and exposed to current or RCP 8.5 climatic conditions for three days. After a one-day incubation period to ambient temperature levels, the soil from the microcosms was collected and poured into a plastic cup (250 mL). The soil moisture was then adjusted to 23% in all treatments. Ten third instar larvae of the root herbivore *D. balteata* were added into each plastic cup. After seven days, all larvae were placed in white traps (White 1927). The parasitism rate and number of emerging juveniles were recorded after 14 days. The number of EPN emerging from larvae infected with EPNs originating from one tube were averaged and used as a proxy for EPN fitness. Testing five strains with sufficient replicate numbers required four independent experiments, each including EN01 and two additional strains (n=7-16 per strain and treatment in total). The infectivity and number of EPNs emerging from the insect cadavers were expressed relatively to EN01-ambient conditions treatment.

### Effects of single climatic conditions on EPN performance and fitness

To appraise the individual and interactive effects of soil temperature, soil moisture, and CO_2_ onto EPN performance and fitness, a full factorial experiment was designed, using all combinations between current and predicted climatic variable levels. The assays were performed using the nematode *H. bacteriophora* strain EN01 and following the same methods as described above. In a first assay, EPN survival was recorded by collecting EPNs using a modified Baermann funnel method (Baermann 1917). Briefly, soil samples were placed into a paper towel and rehydrated until saturation in a funnel, which was sealed with metal clips at the bottom. EPN extraction was performed at ambient temperature conditions. Extracts with suspended EPNs were collected after 24 and 48 hours. The proportion of EPNs collected after 24 hours over the total recovered EPN number was used as a proxy for mobility. In a second assay, EPNs were collected and used for infectivity and fitness assessment as described above. This experiment was repeated three times to ensure good replicate numbers (n=6-12 per treatment).

### Statistical analyses

Statistical analyses were conducted using R (version 3.5.3, https://www.r-project.org) and online tools (http://quantpsy.org; https://www.graphpad.com). Normality and heteroscedasticity of error variance were assessed using Levene’s and Shapiro-Wilk tests, as well as by visualizing quantile-quantile plots and model residuals versus fitted values. The survival rates of spiders and ladybugs were analysed using chi-square tests incorporating a Yates correction for continuity. The survival rate of wasps was analysed using a generalized linear model using the proportion of surviving wasps weighed by the number of wasps in each tube as a response to climatic scenarios. Spider predation rates was analysed using linear models. Ladybug predation and wasp parasitism rates were analysed using generalized linear models with a binomial error structure. EPN infectivity and emergence data were expressed relatively to the EN01 strain in ambient conditions. EPN emergence data from insect infected with EPNs originating from the same microcosm tube were averaged and ln transformed prior analysis. One emergence data point was considered as outlier (Grubbs’ test, p<0.05) and removed from the analysis. EPN relative infectivity and fitness were analysed using linear models. Climatic scenarios/variables, herbivore species (when compared within a single experiment) and nematode strains were treated as fixed effects. Cyclers and CO_2_ channels did not increase the model gain on likelihood and were therefore removed from the models. Experimental repetitions were treated as fixed variables. Significant models were evaluated by Tukey Honest Significant Differences (TukeyHSD) post hoc tests.

## RESULTS

### Predicted climatic conditions do not directly influence the survival of aboveground predators

The designed microcosms ensured good survival rates of wolf spiders, ladybugs and parasitoid wasps over the course of the experiment (Figure 1). Exposure to RCP 8.5 climate conditions, including elevated levels of CO_2_, elevated temperature, and decreased moisture, did not impair the survival of any of the tested arthropods (Figure 1A, C, E).

**Figure 1.**
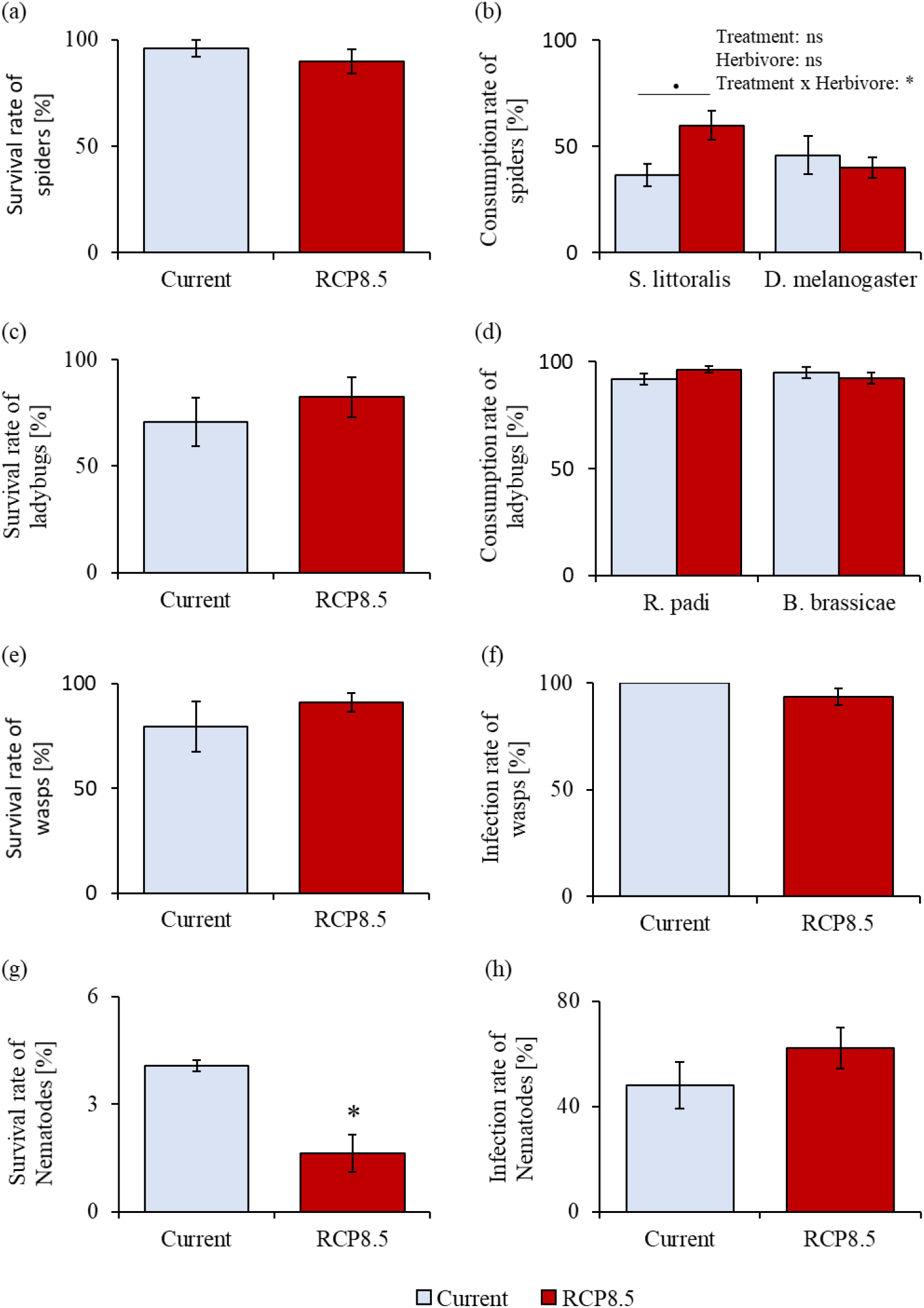
Survival and predation efficacy of herbivore natural enemies after exposure to current and predicted RCP 8.5 climate scenario (IPCC 2014). Survival (Yates’ Chi square, mean ± se, n= 20-24) and consumption rate (linear models, means ± se, n = 9-12) of the wolf spider, *Alopecosa albofasciata*, after exposure to temperature, precipitation and CO_2_ levels of current or predicted (RCP 8.5) climate scenarios (a, b). Survival (Yates’ Chi square, means ± se, n = 17) and consumption rate (generalized linear models, means ± se, n = 6-14) of the ladybird, *Adalia bipunctata*, after exposure to temperature, precipitation and CO_2_ levels of current or predicted (RCP 8.5) climate scenarios (c, d). Survival (generalized linear model, means ± se, n = 5-6) and infection rate (generalized linear models, means ± se, n = 5-6) of the parasite wasp, *Aphidius ervi*, after exposure to temperature, precipitations and CO_2_ levels of current or predicted (RCP 8.5) climate scenarios (e, f). Survival rate (generalized linear model, means ± se, n = 2-3) and infection rate (generalized linear models, means ± se, n = 6-9) of the commercial entomopathogenic nematode, strain EN01, under exposure to temperature, moistures and CO_2_ corresponding to current or predicted (RCP 8.5) climate scenarios (g, h). Stars indicate significant differences, *: p < 0.05. Dots indicate trends, .: 0.05 < p < 0.10.

### Predicted climatic conditions altered aboveground predation in a species-specific manner

Exposure to current or predicted RCP 8.5 climatic conditions resulted in species-specific changes in predation success. The combination of elevated CO_2_, increased temperature, and decreased moisture (RCP 8.5) altered the spider predation rates in a prey-specific manner. It slightly increased spider consumption of the cotton leafworm, *S. littoralis*, but not of the fly larvae, *D. melanogaster* (Figure 1B). Exposure of ladybugs to current or predicted RCP 8.5 climatic conditions resulted in similar consumption rates of cabbage- and bird cherry-oat aphids (Figure 1D). Similarly, exposure to current or predicted RCP 8.5 climatic conditions did not alter the wasp parasitism rate of cabbage aphids (Figure 1F).

### Predicted climatic conditions increase herbivore infection rates by nematodes

Direct exposure to current or predicted RCP 8.5 climatic conditions in microcosms did not decrease infection rates across five tested EPN strains (Figure 2A). On average, exposure to RCP 8.5 conditions increased EPN infection rates by 10.03 %. The effect of the RCP 8.5 scenario was the strongest on EN01, whose infection rates increased by 21.44 % compared to current conditions. Exposure to predicted RCP 8.5 scenarios did not alter the number of emerging juveniles per insect host in any of the tested strains, nor the overall progeny number (Figures 2B-C).

**Figure 2.**
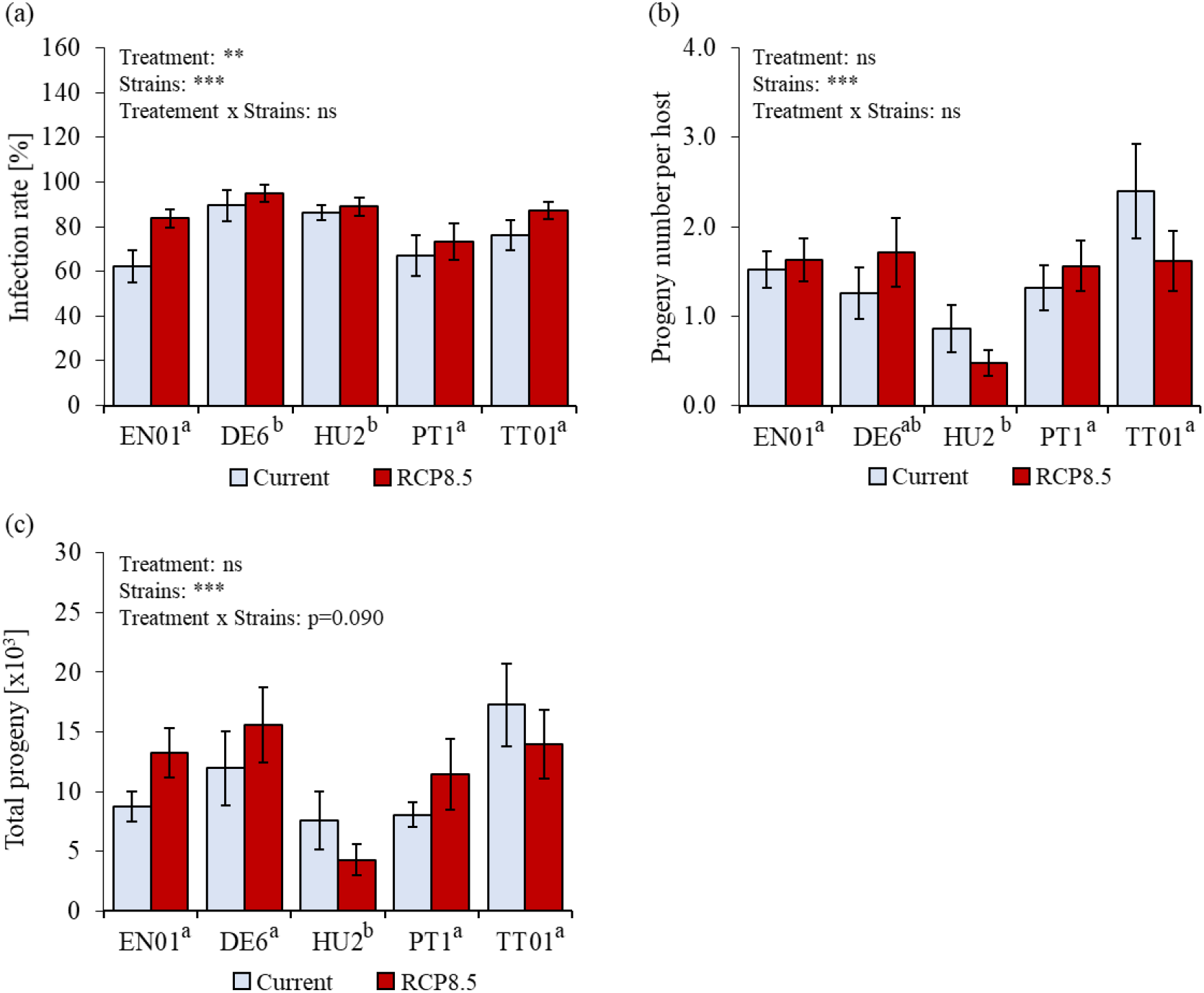
Entomopathogenic nematode infection rate and progeny numbers after exposure to current and predicted RCP 8.5 climate scenario (IPCC 2014). Infection rate (generalized linear model, mean ± se, n = 7-16) of five different strains of the entomopathogenic nematode *H. bacteriophora* on *Diabrotica balteata* larvae after exposure to temperature, precipitations and CO_2_ corresponding to current or predicted (RCP 8.5) climate scenarios (a). Number of infective juveniles (generalized linear model, mean ± se, n = 6-16) emerging per infected host cadaver in five strains (b). Total progeny number calculated as the product between the number of infective juveniles emerging per host and the number of infected hosts (generalized linear model, mean ± se, n = 6-12) (c). Stars indicate significant differences: ***: p ≤ 0.001; **: p ≤ 0.01; ns: non-significant. Different superscript letters next to the strain names indicate significant differences among strains.

### Reduced soil moisture decreases EPN mobility, but increases EPN fitness

Factorial manipulation of the individual climatic variables revealed that exposure to lower soil moisture levels (−28% precipitation) modulated EPN success, while exposure to increased temperature and elevated CO_2_ levels had no additional effects. Reduced soil moisture slightly decreased EPN survival (albeit not significantly), and mobility (Figures 3A-B), but increased infection rates (Figure 3C). While the impact of soil moisture on the progeny number per host was negligible (Figure 3D), the overall EPN fitness was increased (Figure 3E).

**Figure 3.**
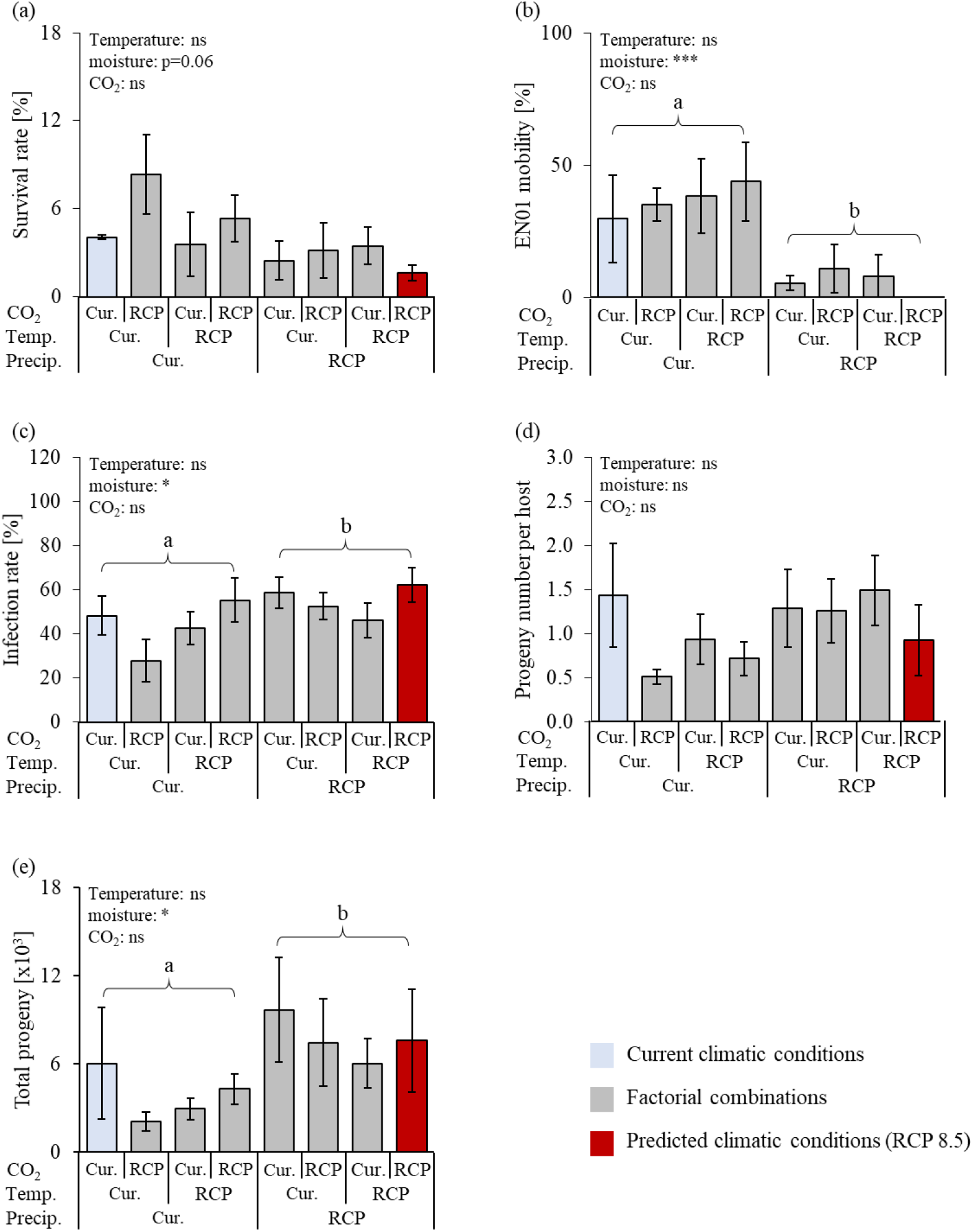
Effects of single and combined climatic variables on entomopathogenic nematode fitness and parasitism efficacy. Survival (linear model, Mean ± se, n = 2-4) of the entomopathogenic nematode (EPN), *Heterorhabditis bacteriophora* (strain EN01), after exposure to current and predicted RCP 8.5 temperature, precipitation, and CO_2_ (a). EPN mobility (generalized linear model, Mean ± se, n = 2-4) after exposure to current and predicted RCP 8.5 temperature, precipitation, and CO_2_ (b). EPN infection rate (generalized linear model, Mean ± se, n = 6-12) of the root herbivore *Diabrotica balteata* larvae after exposure to current and predicted RCP 8.5 temperature, precipitation, and CO_2_ (c). EPN progeny number (IJs) per infected *D. balteata* larva (linear model, Mean ± se, n = 4-11) after exposure to current and predicted RCP 8.5 temperature, precipitation, and CO_2_ (d). EPN total progeny (generalized linear model, Mean ± se, n= 4-11) after exposure to current and predicted RCP 8.5 temperature, precipitation, and CO_2_ (e). Temp.: Temperature; Precip.: Precipitation; Cur.: Current, RCP: RCP 8.5 (IPCC 2014). Stars indicate significant differences: ***: p ≤ 0.001; *: p ≤ 0.05; ns: non-significant. Different letters indicate significant differences. No interaction between the climatic variables was found to be significant.

## DISCUSSION

Our study highlights that a short exposure to temperature, CO_2_ and precipitation levels predicted under an extreme climatic scenario (RCP 8.5), does not impair the efficacy of different natural enemies of herbivores, including several important biological control agents. We even observed a prey species-specific increase in spider efficiency and enhanced nematode infection rates. The possible underlying mechanisms and ecological relevance of these observations are discussed below.

Current models predict that climate change will strongly harm higher trophic levels (Voigt *et al*. 2003; Thakur 2020). Our study reveals that short-term climatic-related changes have negligible direct effects on the ability of third trophic level organisms to predate or parasitize herbivores. This finding suggests that the negative impact of climate change on higher trophic levels observed in large meta-analyses and field experiments may mostly be mediated through either long-term exposure or indirect (plant and herbivore-mediated) changes. Exposure to future climatic scenario did not affect the survival of aboveground herbivore natural enemies such as spiders, ladybugs and parasitoid wasps but decreased the survival of belowground entomopathogenic nematodes (EPNs) within four days. This observation concurs with the hypothesis that arthropods, possessing a waxy cuticle that protects them from desiccation, may be more resistant to climate change than soft-bodied organisms (Sylvain *et al*. 2014). While arthropod predators and parasites are prevalent aboveground (Nyffeler & Birkhofer 2017), EPNs are key herbivore enemies belowground (Půza & Mrácek 2005; Denno *et al*. 2008). The contrasting impact of climate change on above- and belowground organisms survival supports the hypothesis that green, aboveground, food webs may be more sensitive to climate change than brown, belowground ones (Thakur 2020). Yet, future climatic conditions did not negatively impact the predation or parasitism rates by herbivore natural enemies, and even increased the success of nematodes and spiders. The increased infection rate by nematodes was sufficient to compensate for their lower survival in term of progeny number. While climate change may not directly impair EPN ability to control herbivore pest population, changes in host abundance, phenology and quality can drastically modulate their efficacy (Půza & Mrácek 2005; Hiltpold *et al*. 2016; Hiltpold *et al*. 2020; Guyer *et al*. 2018). Interestingly, the increase in spider predation rate was prey species-specific. Such shift in prey consumption was previously observed in Arctic spiders, which ultimately led to slower decomposition rates in soil (Koltz *et al*. 2018). As spiders consume up to 800 million tons of prey per year worldwide (Nyffeler & Birkhofer 2017), identifying future shifts in their predation strategy will be crucial to reliably predict changes in food webs and ecosystem functioning.

Understanding how different climatic variables act together to impact the efficiency of natural enemies of herbivores is an important frontier in multitrophic interaction ecology. Investigating the impact of single and combined climate related drivers onto EPN survival and efficacy revealed no interaction between elevated temperature, CO_2_, and decreased precipitation. While temperature is known to be a major driver of EPN survival and infectivity (Pervez *et al*. 2015; Lalramliana & Yadav 2016; Aatif *et al*. 2020), we did not find any impact of warming in our study. This discrepancy may be explained by the low range of temperature used in our study (19.6 - 23°C) compared to the described beneficial temperature range (25 - 30 °C) (Pervez *et al*. 2015). Consistently with previous literature, elevated levels of CO_2_ did not affect EPN survival and parasitism (Hiltpold *et al*. 2020). Instead, lower precipitation alone seemed to contribute to increased infection rates and progeny number, while decreasing the nematode survival and mobility in the soil. Nematodes have evolved physiological and behavioural strategies to cope with low environmental moisture (Grewal *et al*. 2011; Kagimu *et al*. 2017). They can respond to desiccation by initiating anhydrobiosis, a phenomenon largely associated with the accumulation of trehalose and water stress-related proteins (Grewal *et al*. 2011; Kagimu *et al*. 2017; Womersley 1990). The tolerance to desiccation conferred by anhydrobiosis varies considerably between EPN species and strains (Grewal *et al*. 2002; Mukuka *et al*. 2008). Furthermore, EPNs can avoid desiccation through aggregative movement patterns (Ruan *et al*. 2018) and migration to deeper soil layers where moisture levels are higher (Salame & Glazer 2015). It was also suggested that EPNs can avoid desiccation by remaining longer inside their hosts (Brown & Gaugler 1997; Koppenhöfer *et al*. 1997; Půza & Mrácek 2005, 2007). It is tempting to speculate that the higher infection rates and progeny numbers observed in our study upon predicted climatic scenario are the result of some EPN desiccation tolerance strategy.

Climate change is a multifactorial phenomenon that may affect multitrophic interactions through direct and indirect effects. Interactive effects of the global change drivers and cascading effects among food webs, render predictions about the response of multitrophic challenging to draw. Our study complements current models and shows that combined elevated temperature, CO_2_ levels and decreased precipitation do not have a direct negative impact on the performance of third trophic level organisms upon short term exposure. Together with the current literature, this suggest that indirect, bottom-up-mediated, climate change effects may be stronger than direct effects on predator and parasite efficacy in controlling herbivore populations. Yet, our study highlighted possible shifts in prey species, while the latter were kept in ambient conditions. This phenomenon may have substantial repercussions on ecosystem functioning and would deserve more attention. Building reliable models of food web response to climate change will be of crucial importance to face future natural and agricultural challenges.

## ACKNOWLEDGEMENTS

We are grateful to Anita Streit, Kristýna Filipová and Elina Vogiatzaki for rearing insects and nematodes. We also thank Martin Tschanz and Remo Diethelm for their technical support while developing the microcosms. We thank Tobias Züst for his expertise with statistical analyses and for comments on a previous version of this manuscript. This work was supported by the University of Bern (UniBe 2021).

## AUTHOR CONTRIBUTIONS

CAMR developed and supervised the project. CvD, CM, XZ performed the experiments. MP built the system. CAMR, CvD, and MP analysed the data. CvD and CAMR wrote the first draft of the manuscript.

## SUPPLEMENTARY FIGURES

**Figure S1.**
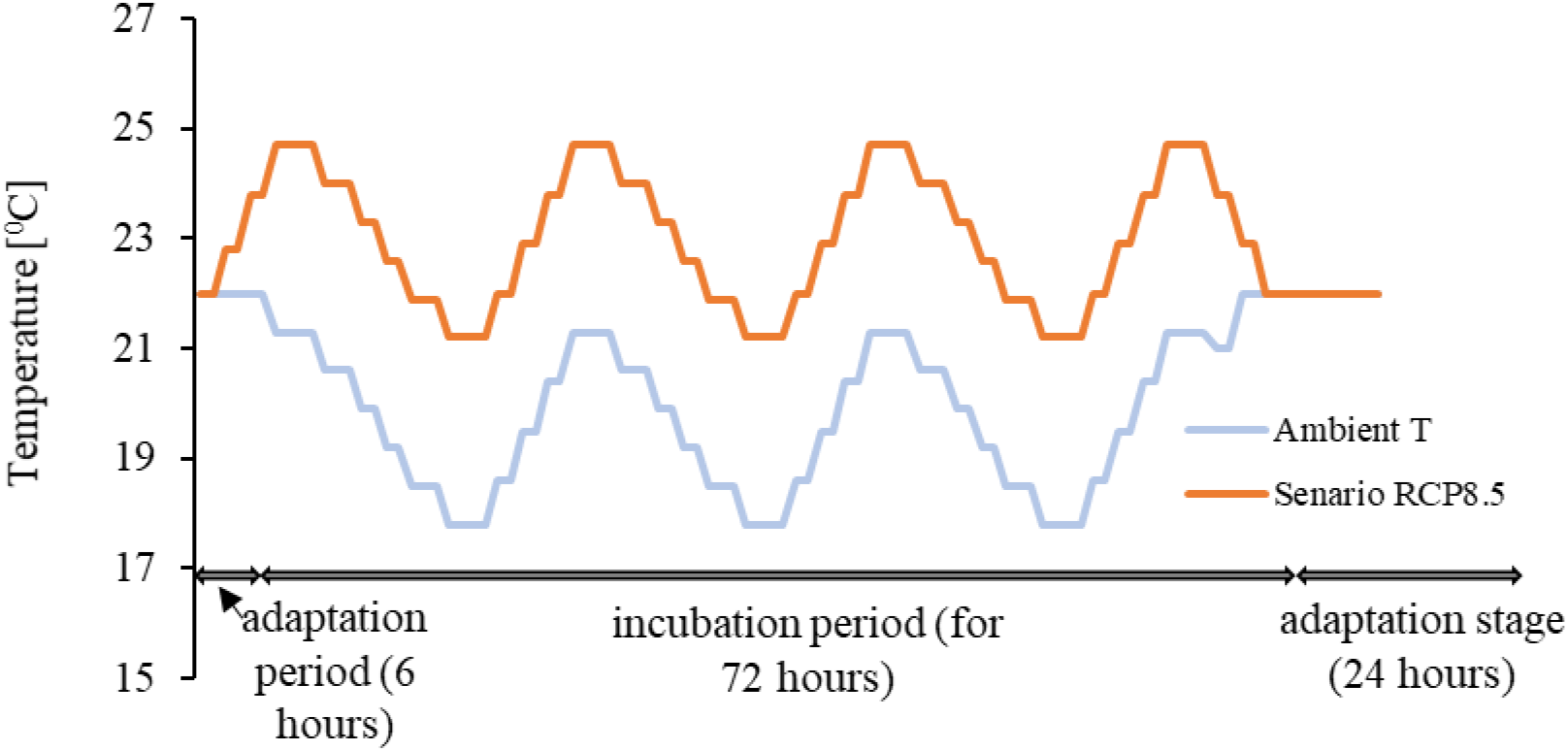
Microcosm temperature regimes. To mimic natural conditions, the two temperature conditions were set to reach a maximum at 4 pm and progressively reduced by 3.5°C to reach a minimum around 6 am. Adaptation stage was set 6 hours before three day-night cycles, after three day-night cycles EPNs equilibrate to 22°C for 16 hours prior to infectivity tests.

**Figure S2.**
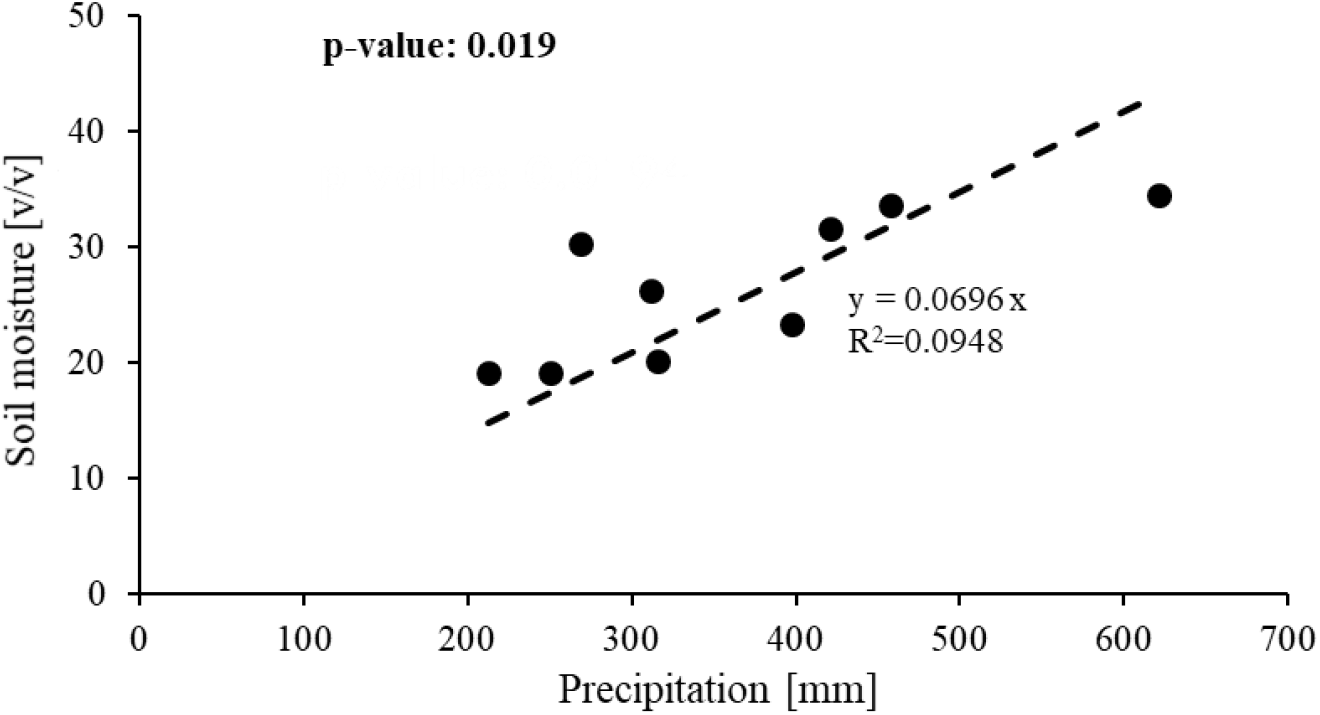
Correlation between precipitation and soil moisture. Linear correlation between average June precipitation sum (in mm) and average June soil moisture (volume/volume) at a soil depth of 15 cm between 2004 and 2016 (years 2005, 2006, 2012 and 2013 were not recorded in the field) in the Swiss Central Plateau (47°17’11.1” N / 7°44’01.5” E), Switzerland.

## REFERENCES

Aatif HM, Hanif MS, Raheel M et al. (2020) Temperature dependent virulence of the entomopathogenic nematodes against immatures of the oriental fruit fly, Bactrocera dorsalis Hendel (Diptera: Tephritidae). Egyptian Journal of Biological Pest Control, 30.

Abdala-Roberts L, Puentes A, Finke DL et al. (2019) Tri-trophic interactions: bridging species, communities and ecosystems. Ecology letters, 22, 2151–2167.

Altieri AH, Bertness MD, Coverdale TC, Herrmann NC, Angelini C (2012) A trophic cascade triggers collapse of a salt-marsh ecosystem with intensive recreational fishing. Ecology, 93, 1402–1410.

Baermann G (1917) A simple method for the detection of *Ankylostomum* (nematode) larvae in soil tests. Geneeskundig Laboratorium te Weltevreden, 57, 131–137.

Barnett KL, Facey SL (2016) Grasslands, Invertebrates, and Precipitation: A Review of the Effects of Climate Change. Frontiers in plant science, 7, 1196.

Bellard C, Bertelsmeier C, Leadley P, Thuiller W, Courchamp F (2012) Impacts of climate change on the future of biodiversity. Ecology letters, 15, 365–377.

Berridge M (2012) Osmoregulation in terrestrial arthropods. Chemical zoology, 5, 287–320.

Bohan DA, Raybould A, Mulder C et al. (2013) Networking agroecology. In: Ecological networks in an agricultural world (eds Woodward G, Bohan DA), pp 1–67. Academic Press, Amsterdam.

Boukal DS, Bideault A, Carreira BM, Sentis A (2019) Species interactions under climate change: connecting kinetic effects of temperature on individuals to community dynamics. Current opinion in insect science, 35, 88–95.

Brown IM, Gaugler R (1997) Temperature and humidity influence emergence and survival of entomopathogenic nematodes. Nematology, 43, 363–375.

Chidawanyika F, Mudavanhu P, Nyamukondiwa C (2019) Global climate change as a driver of bottom-up and top-down factors in agricultural landscapes and the fate of host-parasitoid interactions. Frontiers in Ecology and Evolution, 7, 80. https://www.frontiersin.org/article/10.3389/fevo.2019.00080.

Cornelissen T (2011) Climate change and its effects on terrestrial insects and herbivory patterns. Neotropical entomology, 40, 155–163.

Crain CM, Kroeker K, Halpern BS (2008) Interactive and cumulative effects of multiple human stressors in marine systems. Ecology letters, 11, 1304–1315.

Damien M, Tougeron K (2019) Prey-predator phenological mismatch under climate change. Current opinion in insect science, 35, 60–68.

Darling ES, Côté IM (2008) Quantifying the evidence for ecological synergies. Ecology letters, 11, 1278–1286.

Denno RF, Gruner DS, Kaplan I (2008) Potential for entomopathogenic nematodes in biological control: a meta-analytical synthesis and insights from trophic cascade theory. Journal of Nematology, 40, 61–72.

Draper AM, Weissburg MJ (2019) Impacts of global warming and elevated co2 on sensory behavior in predator-prey interactions: a review and synthesis. Frontiers in Ecology and Evolution, 7.

Furlong MJ, Zalucki MP (2017) Climate change and biological control: the consequences of increasing temperatures on host-parasitoid interactions. Current opinion in insect science, 20, 39–44.

Gravel D, Albouy C, Thuiller W (2016) The meaning of functional trait composition of food webs for ecosystem functioning. Philosophical Transactions of the Royal Society of London. Series B, Biological sciences, 371.

Grewal PS, Bai XD, Jagdale GB (2011) Longevity and stress tolerance of entomopathogenic nematodes. In: Molecular and physiological basis of nematode survival (eds Perry RN, Wharton DA), pp 157–181. CABI, Wallingford.

Grewal PS, Wang X, Taylor RAJ (2002) Dauer juvenile longevity and stress tolerance in natural populations of entomopathogenic nematodes: is there a relationship? International Journal for Parasitology, 32, 717–725.

Guyer A, Hibbard BE, Holzkämper A, Erb M, Robert CA (2018) Influence of drought on plant performance through changes in belowground tritrophic interactions. Ecology and evolution, 8, 6756–6765.

Han P, Becker C, Sentis A, Rostás M, Desneux N, Lavoir A-V (2019) Global change-driven modulation of bottom-up forces and cascading effects on biocontrol services. Current opinion in insect science, 35, 27–33.

Hance T, van Baaren J, Vernon P, Boivin G (2007) Impact of extreme temperatures on parasitoids in a climate change perspective. Annual Review of Entomology, 52, 107–126.

Hiltpold I, Johnson SN, Le Bayon R-C, Nielsen UN (2016) Climate change in the underworld: impacts for soil-dwelling invertebrates. In: Global Climate Change and Terrestrial Invertebrates (eds Johnson SN, Jones TH), pp 201–228. John Wiley & Sons, Ltd.

Hiltpold I, Moore BD, Johnson SN (2020) Elevated atmospheric carbon dioxide concentrations alter root morphology and reduce the effectiveness of entomopathogenic nematodes. Plant and Soil, 447, 29–38. https://doi.org/10.1007/s11104-019-04075-0.

Hu L, Robert CA, Cadot S et al. (2018) Root exudate metabolites drive plant-soil feedbacks on growth and defense by shaping the rhizosphere microbiota. Nature Communications, 9, 2738.

IPCC (2014) Climate change 2014: synthesis report. contribution of working groups i, ii and iii to the fifth assessment report of the intergovernmental panel on climate change [Core Writing Team, R.K. Pachauri and L.A. Meyer (eds.)]. Intergovernmental panel on climate change [Stocker, T.F., D. Qin, G.-K. Plattner, M. Tignor, S.K. Allen, J. Boschung, A. Nauels, Y. Xia, V. Bex and P.M. Midgley (eds.)].

Jactel H, Koricheva J, Castagneyrol B (2019) Responses of forest insect pests to climate change: not so simple. Current opinion in insect science, 35, 103–108.

Jamieson MA, Trowbridge AM, Raffa KF, Lindroth RL (2012) Consequences of climate warming and altered precipitation patterns for plant-insect and multitrophic interactions. Plant Physiology, 160, 1719–1727. https://pubmed.ncbi.nlm.nih.gov/23043082.

Jeffs CT, Lewis OT (2013) Effects of climate warming on host–parasitoid interactions. Ecological Entomology, 38, 209–218.

Jun Chen F, Wu G, Parajulee MN, Ge F (2007) Impact of elevated CO2 on the third trophic level: A predator *Harmonia axyridis* and a parasitoid Aphidius picipes. Biocontrol Science and Technology, 17, 313–324.

Kagimu N, Ferreira T, Malan AP (2017) The attributes of survival in the formulation of entomopathogenic nematodes utilised as insect biocontrol agents. African Entomology, 25, 275–291.

Kaplan I, Carrillo J, Garvey M, Ode PJ (2016) Indirect plant-parasitoid interactions mediated by changes in herbivore physiology. Current opinion in insect science, 14, 112–119.

Kharouba HM, Ehrlén J, Gelman A, Bolmgren K, Allen JM, Travers SE, Wolkovich EM (2018) Global shifts in the phenological synchrony of species interactions over recent decades. Proceedings of the National Academy of Sciences, 115, 5211–5216.

Koltz AM, Classen AT, Wright JP (2018) Warming reverses top-down effects of predators on belowground ecosystem function in Arctic tundra. Proceedings of the National Academy of Sciences, 115, E7541–E7549. https://www.pnas.org/content/115/32/E7541.

Koppenhöfer AM, Baur ME, Stock SP, Choo HY, Chinnasri B, Kaya HK (1997) Survival of entomopathogenic nematodes within host cadavers in dry soil. Applied Soil Ecology, 6, 231–240.

Kreyling J, Beier C (2013) Complexity in climate change manipulation experiments. BioScience, 63, 763–767. www.jstor.org/stable/10.1525/bio.2013.63.9.12.

Lalramliana, Yadav AK (2016) Effects of storage temperature on survival and infectivity of three indigenous entomopathogenic nematodes strains (Steinernematidae and Heterorhabditidae) from Meghalaya, India. Journal of parasitic diseases: official organ of the Indian Society for Parasitology, 40, 1150–1154.

Laws AN (2017) Climate change effects on predator-prey interactions. Current opinion in insect science, 23, 28–34.

Majeed S, Hill SR, Ignell R (2014) Impact of elevated CO2 background levels on the host-seeking behaviour of *Aedes aegypti*. The Journal of experimental biology, 217, 598–604.

McMullen II JG, Stock SP (2014) *In vivo* and *in vitro* rearing of entomopathogenic nematodes (Steinernematidae and Heterorhabditidae). JoVE, e52096. https://www.jove.com/video/52096.

Melguizo-Ruiz N, Jiménez-Navarro G, Moya-Laraño J (2016) Beech cupules as keystone structures for soil fauna. PeerJ, 4, e2562.

Mukuka J, Strauch O, Ehlers RU (2008) Variability in desiccation tolerance among different strains of the entomopathogenic nematodes *Heterorhabditis bacteriophora*. Communications in Agricultural and Applied Biological Sciences, 73, 669–672.

Newman JA, Anand M, Henry HAL, and Hunt SL (2011) Climate change biology. CABI, Wallingford, Oxon., Cambridge, Mass.

Nyffeler M, Birkhofer K (2017) An estimated 400-800 million tons of prey are annually killed by the global spider community. Die Naturwissenschaften, 104, 30.

Parmesan C, Yohe G (2003) A globally coherent fingerprint of climate change impacts across natural systems. Nature, 421, 37–42.

Pervez R, Eapen SJ, Devasahayam S, Jacob TK (2015) Effect of temperature on the infectivity of entomopathogenic nematodes against shoot borer (*Conogethes punctiferalis* Guen.) infesting ginger (*Zingiber officinale* Rosc.). Journal of Biological Control, 29, 187–193.

Pincebourde S, van Baaren J, Rasmann S, Rasmont P, Rodet G, Martinet B, Calatayud P-A (2017) Plant–insect interactions in a changing world. In: Insect-plant interactions in a crop protection perspective (eds Sauvion N, Calatayud P-A, Thiéry D), pp 289–332. Academic Press.

Půza V, Mrácek Z (2005) Seasonal dynamics of entomopathogenic nematodes of the genera *Steinernema* and *Heterorhabditis* as a response to abiotic factors and abundance of insect hosts. Journal of invertebrate pathology, 89, 116–122.

Půza V, Mrácek Z (2007) Natural population dynamics of entomopathogenic nematode *Steinernema affine* (Steinernematidae) under dry conditions: Possible nematode persistence within host cadavers? Journal of invertebrate pathology, 96, 89–92.

Renner SS, Zohner CM (2018) Climate change and phenological mismatch in trophic interactions among plants, insects, and vertebrates. Annual Review of Ecology, Evolution, and Systematics, 49, 165–182.

Robinet C, Roques A (2010) Direct impacts of recent climate warming on insect populations. Integrative zoology, 5, 132–142.

Robinson EA, Ryan GD, Newman JA (2012) A meta-analytical review of the effects of elevated CO2 on plant-arthropod interactions highlights the importance of interacting environmental and biological variables. The New phytologist, 194, 321–336.

Romo CM, Tylianakis JM (2013) Elevated temperature and drought interact to reduce parasitoid effectiveness in suppressing hosts. PloS one, 8, e58136.

Rosenblatt AE, Schmitz OJ (2014) Interactive effects of multiple climate change variables on trophic interactions: a meta-analysis. Climate Change Responses, 1.

Rosenblatt AE, Schmitz OJ (2016) Climate change, nutrition, and bottom-up and top-down food web processes. Trends in Ecology & Evolution, 31, 965–975.

Ruan W-B, Shapiro-Ilan D, Lewis EE, Kaplan F, Alborn H, Gu X-H, Schliekelman P (2018) Movement patterns in Entomopathogenic nematodes: Continuous *vs*. temporal. Journal of invertebrate pathology, 151, 137–143.

Salame L, Glazer I (2015) Stress avoidance: vertical movement of entomopathogenic nematodes in response to soil moisture gradient. Phytoparasitica, 43, 647–655.

Sassi C de, Tylianakis JM (2012) Climate change disproportionately increases herbivore over plant or parasitoid biomass. PloS one, 7, e40557.

Selvaraj P, Ganeshamoorthi P, Pandiaraj T (2013) Potential impacts of recent climate change on biological control agents in agro-ecosystem: A review. international journal of biodiversity and conservation, 5, 845–852. DOI: 10.5897/IJBC2013.0551.

Sylvain ZA, Wall DH, Cherwin KL, Peters DP, Reichmann LG, Sala OE (2014) Soil animal responses to moisture availability are largely scale, not ecosystem dependent: insight from a cross-site study. Global Change Biology, 20, 2631–2643.

Thakur MP (2020) Climate warming and trophic mismatches in terrestrial ecosystems: the green–brown imbalance hypothesis. Biology Letters, 16, 20190770.

Thomson LJ, Macfadyen S, Hoffmann AA (2010) Predicting the effects of climate change on natural enemies of agricultural pests. Biological Control, 52, 296–306.

Torode MD, Barnett KL, Facey SL, Nielsen UN, Power SA, Johnson SN (2016) Altered precipitation impacts on above- and below-ground grassland invertebrates: summer drought leads to outbreaks in spring. Frontiers in Plant Science, 7, 1468.

Tylianakis JM, Didham RK, Bascompte J, Wardle DA (2008) Global change and species interactions in terrestrial ecosystems. Ecology letters, 11, 1351–1363.

van der Putten WH, Macel M, Visser ME (2010) Predicting species distribution and abundance responses to climate change: why it is essential to include biotic interactions across trophic levels. Philosophical transactions of the Royal Society of London. Series B, Biological sciences, 365, 2025–2034.

van der Putten WH, Ruiter PC de, Martijn Bezemer T, Harvey JA, Wassen M, Wolters V (2004) Trophic interactions in a changing world. Basic and Applied Ecology, 5, 487–494.

Vidal MC, Murphy SM (2018) Bottom-up vs. top-down effects on terrestrial insect herbivores: a meta-analysis. Ecology letters, 21, 138–150.

Voigt W, Perner J, Davis AJ et al. (2003) Trophic levels are differentially sensitive to climate. Ecology, 84, 2444–2453.

Walther G-R (2010) Community and ecosystem responses to recent climate change. Philosophical Transactions of the Royal Society B: Biological Sciences, 365, 2019–2024.

Walther G-R, Post E, Convey P et al. (2002) Ecological responses to recent climate change. Nature, 416, 389–395.

White GF (1927) A method for obtaining infective nematode larvae from cultures. Science, 66, 302–303. https://science.sciencemag.org/content/66/1709/302.2.

Willmer PG (1982) Microclimate and the environmental physiology of insects. In: Advances in insect physiology (eds Berridge MJ, Treherne JE, Wigglesworth VB), pp 1–57. Academic Press, London, New York.

Womersley CZ (1990) Dehydration survival and anhydrobiotic potential. In: Entomopathogenic nematodes in biological control, pp 117–137. CRC Press Boca Raton, FL.

Woodward, G, Bohan, DA (Eds.) (2013) Ecological networks in an agricultural world. Academic Press, Amsterdam.

Zhang X, van Doan C, Arce CC et al. (2019) Plant defense resistance in natural enemies of a specialist insect herbivore. Proceedings of the National Academy of Sciences, 116, 23174–23181. https://www.pnas.org/content/116/46/23174.

